# The Posterior Insular Cortex is Necessary for the Consolidation of Tone Fear Conditioning

**DOI:** 10.1101/2020.08.12.248294

**Authors:** J.P.Q. de Paiva, A.P.A. Bueno, M. Dos Santos Corrêa, M.G.M. Oliveira, T.L Ferreira, R.V. Fornari

## Abstract

The insular cortex (IC) is notably implicated in emotional and cognitive processing; however, little is known regarding to what extent its two main subregions play functionally distinct roles on memory consolidation of conditioned fear tasks. Here we verified the effects of temporary functional inactivation of the anterior (aIC) and posterior IC (pIC) on contextual and tone fear memory. Rats received post-training bilateral infusions of the GABA_A_ receptor agonist muscimol into either the aIC or pIC and were tested 48 and 72 hours after the conditioning session to assess contextual (CFC) and tone (TFC) fear conditioning, respectively. Inactivation of the aIC during memory consolidation did not affect fear memory for CFC or TFC. On the other hand, post-training inactivation of the pIC impaired TFC but not CFC. Our findings indicate that the pIC is a necessary part of the neural circuitry related to the consolidation of cued-fear memories.

**Highlights:**

- We studied the role of the anterior (aIC) and posterior (pIC) insula in fear memory
- Post-training inactivation of aIC and pIC did not impact contextual fear conditioning
- The pIC but not aIC is necessary for the consolidation of tone fear conditioning

## 1. INTRODUCTION

The insula (or insular cortex, IC) has been implicated in a myriad of functions such as emotion regulation, cognition, interoception, homeostasis and salience processing (Allen, 2020; Benarroch, 2019; Gogolla, 2017; Nieuwenhuys, 2012; Uddin, 2015; Uddin et al., 2014, 2017). Animal studies have further revealed a key role for the IC in aversive states (Gehrlach et al., 2019; Méndez-Ruette et al., 2019), multimodal processing and integration of visceral and somatosensory inputs (Livneh et al., 2017, 2020; Rodgers et al., 2008).

Traditionally, two major IC subdivisions - an anterior (rostral, aIC) and a posterior (caudal, pIC) - have been described based on their thalamic and amygdalar connections (McDonald, 1998; McDonald et al., 1999; Nieuwenhuys, 2012; Shi & Cassell, 1998b, 1998a; van der Kooy et al., 1984). Recently, a comprehensive viral tracing study in mice has also highlighted the existence of two IC clusters characterized by a markedly distinct connectivity profile along the antero-posterior IC axis (Gehrlach et al., 2020). The different cytoarchitectural and connectivity patterns of the aIC and pIC warrant further studies on a possible functional dissociation at the IC subregion level (Gehrlach et al., 2020; Gogolla, 2017). Some evidences suggest a functional heterogeneity of the IC regarding anxiety (Méndez-Ruette et al., 2019) and some memory processes, such as the acquisition of conditioned taste aversion and spatial memory (Nerad et al., 1996). Nevertheless, there is still considerable debate regarding the role of the aIC and pIC in fear memory.

Initial investigations into the role of the aIC in memory have focused primarily in recognition memory tasks (Bermudez-Rattoni, 2014), but also in the spatial version of the Morris water maze (Bermudez-Rattoni et al., 1991; Nerad et al., 1996) and the inhibitory avoidance (Bermúdez-Rattoni et al., 1997; Fornari, Wichmann, Atucha, et al., 2012; Miranda & Bermúdez-Rattoni, 2007; Miranda & McGaugh, 2004). Nevertheless, the precise role of the aIC in fear conditioning is still inconclusive. For example, pre-training excitotoxic lesions of the more rostral portion of the aIC impaired the acquisition of Contextual Fear Conditioning (CFC), without interfering with Tone Fear Conditioning (TFC, Morgan & LeDoux, 1995). However, temporary synaptic inhibition of this same area attenuated the conditioned freezing response to the context when performed immediately after conditioning or before testing, but not before training, suggesting an effect in consolidation and expression but not in acquisition of CFC (Alves et al., 2013).

Regarding the pIC, a recent study reported that, although a subset of pIC neurons were activated in response to tail shocks, the optogenetic inactivation of this area did not affect the freezing response to footshocks during the fear conditioning training session, nor during the contextual or tone fear recall (Gehrlach et al., 2019). Previous studies have also shown that pre-training lesions of the pIC did not prevent the acquisition of conditioned freezing responses to contextual or tone stimuli (Brunzell & Kim, 2001; Lanuza et al., 2004). However, pre-training reversible inactivation or post-training protein synthesis inhibition of the pIC disrupted the freezing response to the tone, measured 24 hours after training, without affecting fear expression during the conditioning session (Casanova et al., 2016). On the other hand, post-training lesions of the pIC have showed either a selective impairment of TFC and no effect in CFC (Brunzell & Kim, 2001), or have impaired conditioned freezing responses to both context and tone stimuli (Corodimas & LeDoux, 1995). In addition to the divergent results, the use of permanent lesions in previous works did not allow discriminating the role of the pIC in consolidation or retrieval processes of both tasks, or even in the expression of the fear response.

The present study aimed to investigate if the IC is necessary for memory consolidation of CFC and TFC tasks and, more precisely, whether there is a functional dissociation between the aIC and pIC on these two tasks, which are knowingly mediated and encoded by distinct neural circuitries (Fanselow, 2010; Ferreira et al., 2003, 2008; LeDoux, 2000; Phillips & LeDoux, 1992). Rats were submitted to a fear conditioning training session, in which a tone was associated with a footshock and, immediately after training, received bilateral infusions of muscimol into the aIC or pIC to temporarily inactivate the region during the consolidation period. Contextual and tone fear memories were later assessed in the same animals to verify the specific roles played by the IC subregions in the consolidation of both tasks.

## 2. MATERIALS AND METHODS

### 2.1. Animals

Forty-six 3-months-old male Wistar rats, weighing between 250 to 350 grams at time of training, were obtained from *Instituto Nacional de Farmacologia* (INFAR-UNIFESP; aIC experiment) and from *Centro de Desenvolvimento de Modelos Experimentais para Medicina e Biologia* (CEDEME-UNIFESP; pIC experiment). Rats were housed 4 to 5 per cage (30 cm × 16 cm × 18 cm) and with *ad libitum* access to food and water. The animals, adapted to the vivarium for at least 1 week, were kept in controlled conditions of temperature (22 ± 2 °C) and light/darkness period of 12/12 (light period starting at 7 am).

Training and testing were performed during the light phase of the cycle. All procedures followed guidelines and standards of *Conselho Nacional de Controle de Experimentação Animal* (CONCEA - Brazilian Council of Animal Experimentation) and were approved by the Ethics Committee on Animal Use (CEUA - protocol numbers: UFABC: 031/2013 and UNIFESP: 6896201213). Each experiment was conducted with different groups of animals and efforts were made to minimize the number of animals and their suffering.

### 2.2. Apparatus

Behavioral experiments were conducted in two automated fear conditioning chambers (Med-Associates, Inc., St. Albans, VT), connected to a computer interface enabling video recording, analysis and measurement of the rat’s freezing behavior in real time. The conditioning box was 32 cm wide, 25 cm high and 25 cm deep (VFC-008), surrounded by a sound attenuating chamber (63.5 cm × 35.5 cm × 76 cm, NIR-022SD), and illuminated by a LED light source (Med Associates NIR-100), which provided visible white light spectrum (450-650nm) and invisible near-infrared light (NIR, 940nm). The chamber was also equipped with a speaker (90 dba, 2.0-kHz) and a stainless-steel grid floor. The chamber’s top and front walls were made of transparent polycarbonate. A NIR video camera (VID-CAM-MONO-4 Fire Wire Video Camera) was attached to the front of the sound attenuating chamber, facing the transparent front wall of the conditioning chamber, allowing to record and measure freezing time even with no visible light. The scrambled footshocks were administered through the grid floor of the cage (AC constant current), controlled by an Aversive Stimulator (ENV-414S). A general activity index was derived in real time from the video stream by a computer software (Video Freeze, Version 1.12.0.0, Med-Associates). The software performed real-time video recordings (30 frames per second) using a set threshold level (20 arbitrary units of movement) previously calibrated with the experimenters’ freezing scores (r^2^ = .997) and used in previous studies (Bueno et al., 2017; dos Santos Corrêa et al., 2019; Moreira-Silva et al., 2018).

The training context (Context A) was characterized by the conditioning chamber in its original configuration, left constantly illuminated and cleaned with 30% alcohol diluted in water after each trial. The novel context (Context B) consisted in a modified chamber, where a floor and a white curved sidewall extending across the back wall were inserted, both plexiglas made. Light in the Context B was kept off and the box was cleaned with a 5% acetic acid solution after each trial.

### 2.3. Surgery for cannula implantation

Rats underwent stereotaxic surgery for cannula implantation, as described previously (Fornari et al., 2008; Fornari, Wichmann, Atucha, et al., 2012; Fornari, Wichmann, Atsak, et al., 2012). Briefly, rats were anaesthetized with an intraperitoneal injection of ketamine (90 mg/Kg; Syntec) and xylazine (50 mg/Kg; Vetbrands). The skull was positioned in a stereotaxic frame (Kopf Instruments) and two stainless-steel guide cannulae (15 mm, 23 gauge) were implanted bilaterally with the cannula tips 2.0 mm above the aIC or pIC, according to the following coordinates (Paxinos & Watson, 2007): aIC: anteroposterior, +1.5 mm from Bregma; mediolateral, ±4,0 mm from the midline; dorsoventral, −4.5 mm from skull surface, at an angle of 4 degrees; pIC: anteroposterior, −2.0 mm from Bregma; mediolateral, ± 5,0 mm from the midline; dorsoventral, −5.0 mm from the skull surface, at an angle of 10 degrees. The cannulae were affixed to the skull with two anchoring screws and dental cement. A thermal pad was placed under the animal to maintain its body temperature throughout the surgery and to avoid possible hypothermia induced by anesthetics. Stainless-steel stylets (15 mm long) were inserted into each cannula to maintain unobstructedness and were removed only for drug infusion.

After surgery, rats received a subcutaneous injection of 3 ml of saline to facilitate clearance of drugs and prevent dehydration, then were administered a veterinary pentabiotic (0.2 ml, Fort Dodge Animal Health, intramuscularly) and an analgesic (Ketoprofen, 0.2 ml, intramuscularly) to avoid bacterial infection and pain. Next, animals were housed in heated individual boxes, until they were fully awakened. Finally, the animals were returned to the vivarium in individual transparent cages and placed side by side, to reduce perception of isolation. The animals were allowed at least 7 days of recovery before behavioral procedures. During the postoperative recovery, behavior, posture, weight, evidence of pain, signal of infection in suture or cannula clogging were assessed daily.

### 2.4. Behavioral Procedures

Animals were handled individually during 1 minute for three days, prior to the conditioning training, to accustom them to the infusion procedure and to minimize stress during the session. All experiments took place between 11am and 5pm. Rats were habituated to the behavioral room for at least 90 minutes before training and CFC testing.

#### Training session

rats were individually placed inside the conditioning box (Context A). After 120 seconds of free exploration, a tone (90 dB, 2000 Hz) was delivered for 30 seconds, coterminating with a footshock (0.7 mA, 1 s). Rats were removed 60 seconds after the end of the shock. During the training session, freezing time was recorded by the system either before (2-minutes baseline freezing) and after (1-minute post-shock freezing) the tone– footshock pairing. Immediately after training, each rat received bilateral microinfusions of muscimol (0.5 μg/0.5 μL) or saline in the aIC (experiment 1) or pIC (experiment 2).

#### Shock reactivity

during the training session, the rats’ responses to electric shocks were recorded (shock reactivity) as previously described (Fornari et al., 2008). For each footshock, one of the following scores was given: 3 = jumping, 2 = vocalization, 1 = flinching, 0 = no response.

#### Conditioning tests

the CFC test was performed 48 hours after training and consisted in the re-exposure to Context A for 5 minutes, with no presentation of tone or shock.

After 24 hours, the same animals were submitted to the TFC test in a new context (Context B). To avoid a generalization effect of the CFC memory in the TFC test, animals were individually transported directly from the vivarium to the fear conditioning room, using a different transport box. Each animal was directly placed in Context B for 4 minutes, and at the end of the second and third minutes, the tone stimulus presented in the training session (90 dB, 2000 Hz) was delivered for 30 seconds. Freezing time was continuously measured during the first 2 minutes before the first tone onset (pre-tone) and during the final 2 minutes after the first tone onset (post-tone). A schematic representation of the experimental design is depicted in Figure 2a. During both CFC and TFC tests, the behavioral observer was blind to the rat’s treatment condition.

**Figure 1.**
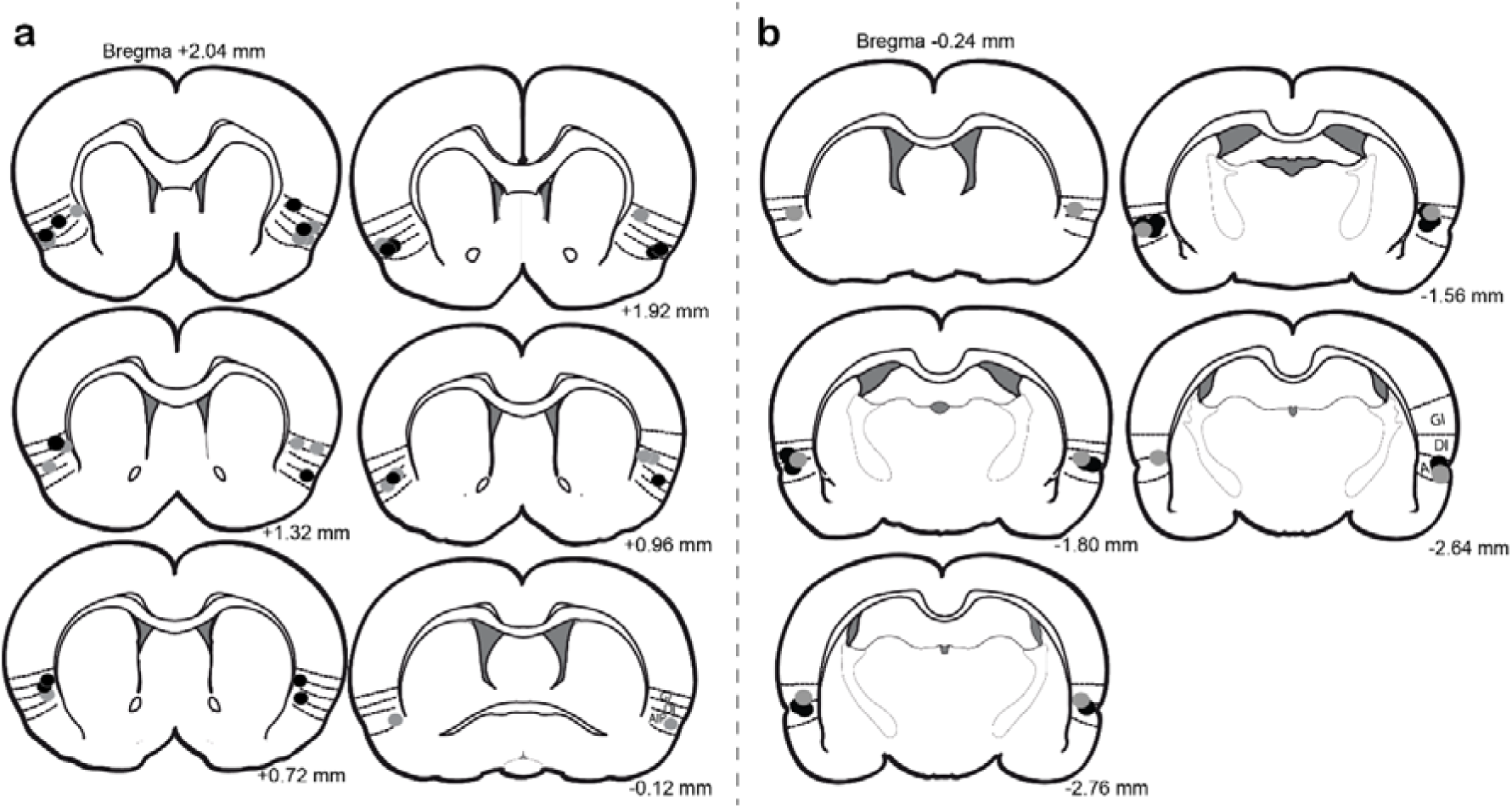
a) Histological verification of microinjection sites for the anterior (aIC) and b) the posterior (pIC) insular cortex. Colored dots represent the center of the needle tip for each rat (gray: saline; black: muscimol). The numbers indicate the distance from the bregma (Paxinos & Watson, 2007).

**Figure 2:**
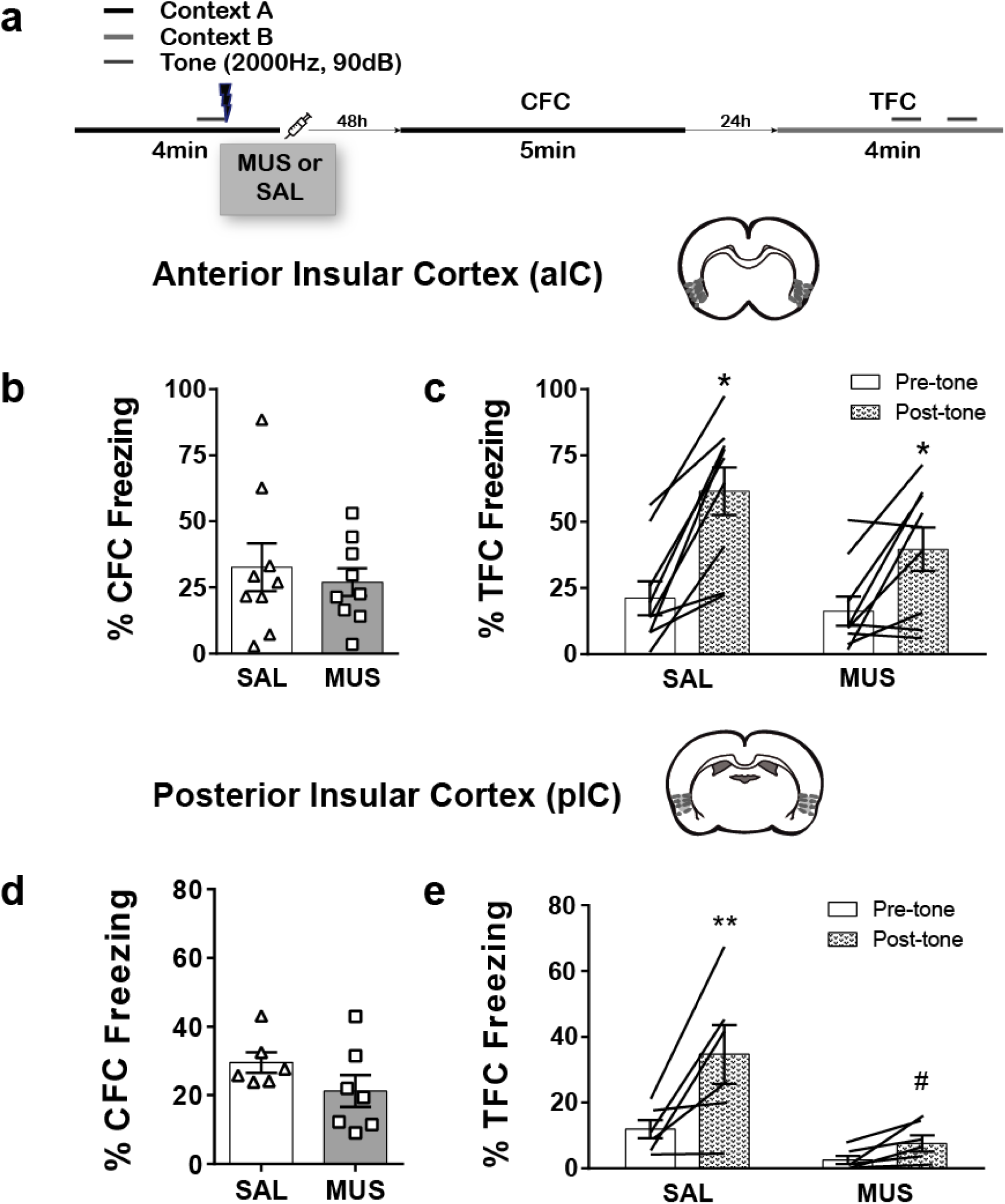
Temporary inactivation of the posterior (pIC) but not the anterior (aIC) insular cortex, during memory consolidation, impairs tone fear conditioning (TFC). (a) Schematic representation of the experimental design; (b,c,d,e) Freezing time percentage (mean ± standard error of the mean) during the contextual fear conditioning (CFC) and TFC test sessions of rats microinjected with saline or muscimol into the aIC (b,c) or pIC (d,e) immediately after training. Triangles represent rats’ individual data for CFC and lines for TFC. The patterned bars represent post-tone freezing times during TFC. (*) p ≤ 0.05, compared to the correspondent group before tone presentations. (**) p < 0.01, compared to the correspondent group before tone presentations. (#) p < 0.01 compared to the control group after tone presentations. CFC: contextual fear conditioning; TFC: tone fear conditioning.

### 2.5. Drug and infusion procedure

Muscimol (Sigma-Aldrich®), a selective GABA_A_ agonist, was dissolved in sterile saline solution (0.9% NaCl). The dose of 0.5 μg/0.5 μL was chosen based on previous studies that successfully inactivated the IC prior to aversive memory conditioning trainings (Berman et al., 2000; Casanova et al., 2016; Foilb et al., 2016; Koh & Bernstein, 2005; Rodríguez et al., 2020).

Immediately after the fear conditioning training, animals were gently immobilized in the experimenter’s lap for bilateral infusions of muscimol (0.5 μL/hemisphere) or an equivalent volume of vehicle (sterile saline) into the targeted IC, by using 30 gauge injection needles connected to a 10 μl Hamilton microsyringe with polyethylene (PE-10) tubing. The injection needle protruded 2.0 mm beyond the tip of the cannula, and the infusions were performed over a period of 60 seconds by an automated syringe pump (Model Bi2000 – *Insight Equipamentos LTDA* Insight, Brazil) at a constant speed (0.5 μL/min). The movement of a small air bubble in the PE-10 tubing before, during and after the infusion confirmed the success of the procedure. The injection needles were kept in place for additional 30 seconds within the cannulae, after drug infusion, to maximize diffusion and to prevent drug backflow. During the microinjection, the animals could move while gently restricted to the lap of the experimenter.

### 2.6 Cannula placement verification

Rats were deeply anesthetized with an overdose of sodium pentobarbital (≈ 100 mg/kg, i.p.) or urethane (25% solution, 2 mL/kg) and then received bilateral infusions of methylene blue dye (1%), at the same volume and speed according to the procedure described previously. Fifteen minutes after the infusion, each rat was decapitated and its brain was removed, frozen over dry ice and stored at −80 ºC. Coronal sections of 40 μm were cut on a cryostat (Leica Microsystems, Germany), mounted on a glass slide and stained with cresyl violet. Sections were taken over a distance considerably rostral and caudal to the insula to assess the infusion site not only within, but also remote to the target area. Mounted slices were examined under a light microscope (Leica Microsystems, model 5500), and the verification of the needle tips injection site into the IC was made according to the atlas of Paxinos & Watson (2007) (Paxinos & Watson, 2007). Composite diagrams have been constructed from these individual representations and are presented in Figure 1. The animals with unclear cannulae location, injection needle placements outside the IC, or with extensive tissue damage were excluded from behavioral analysis.

### 2.7. Statistical Analysis

Fear memory was assessed as the quantity of time the animal spent freezing during the training, CFC and TFC testing sessions. All data were analyzed using JAMOVI (The jamovi Project, 2020). Data are expressed as mean percent freezing times (± standard error of the mean, S.E.M.) and median shock reactivity (upper-lower quartiles). All continuous data were tested for normality using the Kolmogorov-Smirnov normality test. Homogeneity of variances was checked with the Levene’s test after each parametric test.

Data from training and CFC sessions were analyzed using a Student t-test for independent samples. Shock reactivity scores were analyzed using Mann–Whitney U pairwise comparisons to verify between-group differences. Finally, freezing times from the TFC test were analyzed by a repeated-measures analysis of variance (ANOVA) with Treatment as the main factor and pre- and post-tone freezing times as the within-subject variables. The ANOVA was followed, when appropriate, by a Bonferroni post-hoc test to determine the source of the detected significances. For all tests, the level of significance was set at p-value of .05. Effect sizes for the repeated measure ANOVAs were reported as generalized eta-squared (η^2^_G_) when inferential tests resulted in statistically significant effects.

## 2. RESULTS

### 3.1. Histological results

Methylene blue dyes were restricted to the targeted areas. In the first experiment, the infusions were restricted to the aIC between +2.76 and −0.12 mm from the bregma (Figure 1a). In the pIC experiment, the marks of the needle tips comprised between −0.24 and −2.92 mm from the bregma (Figure 1b).

Histological results were used as exclusion criteria. All the animals presenting misplaced injections were excluded from statistical analysis (33% from total). After applying the exclusion criteria, 18 animals (9 muscimol) for the aIC and 13 (7 muscimol) for the pIC were considered for statistical analysis.

### 3.2. Experiment 1: Effects of temporary inactivation of the aIC

During the training session, both groups showed similar freezing times during the first two minutes of exploration before the tone-shock pairing (t(16) = −1.41, p = 0.18). Shock reactivity was also similar for both groups (U = 24.50, p = 0.10). Finally, freezing times at the last minute were also similar between both groups (t(16) = −1.72, p = 0.11). These results are summarized in Table 1.

**Table 1.** Freezing response and shock reactivity of rats during training in both aIC and pIC experiments.

| Group |  | Baseline Freezing<sup>□</sup> | Post-shock Freezing<sup>□</sup> | Shock Reactivity<sup>□</sup> | N |
| --- | --- | --- | --- | --- | --- |
| aIC (Exp 1) | Saline | 4.32 ± 1.39 | 22.54 ± 3.66 | 2 (2-3) | 9 |
|  | Muscimol | 1.92 ± 0.99 | 12.68 ± 4.43 | 2 (2-2) | 9 |
| pIC (Exp 2) | Saline | 0.67 ± 0.79 | 25.51 ± 5.49 | 1.5 (1-2) | 6 |
|  | Muscimol | 1.30 ± 0.24 | 26.05 ± 2.37 | 2 (1.5-2) | 7 |

*Note.* <sup>□</sup>Data are expressed as mean ± standard error of the mean. <sup>□</sup>Data are expressed as median (upper-lower quartile). aIC: anterior insular cortex; pIC: posterior insular cortex.

Two days after training, animals were exposed to the training context but without tone or footshock presentation. The post-training infusion of muscimol did not impair the animals’ freezing times when compared to the saline control group in CFC (Figure 2b, (*t*(16) = −0.54, *p* = 0.59).

Additionally, post-training treatment with muscimol did not impair animals’ freezing time when tested for the TFC a day later (Figure 2c). Repeated-measures ANOVA indicated no effect of Treatment between subjects (*F*(1,16) = 2.13 *p* = 0.16) nor interaction between Treatment and Tone (*F*(1,16) = 2.80, *p* = 0.11, η^2^_G_ = 0.04), but a significant effect of Tone within subjects (*F*(1,16) = 39.48, *p* < 0.001, η^2^_G_ = 0.37).

### 3.3. Experiment 2: Effects of temporary inactivation of the posterior insular cortex

During training, both groups showed similar freezing times during the first two minutes of exploration before tone-shock pairing (t(11) = 0.717, p = 0.49). Shock reactivity was also similar for both groups (U = 19.0, p = 0.81). Freezing times at the last minute were also similar between both groups (t(11) = 0.09, p = 0.93; Table 1).

Two days after training, animals were re-exposed to the training context without tone or footshock, for the CFC test. As shown in Figure 2d, post-training infusion of muscimol did not alter the freezing behavior when compared to the saline group (*t*(11) = −1.44, *p* = 0.18). However, post-training inactivation of the pIC impaired freezing to the tone during the TFC test, on the third day (Figure 2e). Repeated-measures ANOVA indicated significant effects for Treatment (*F*(1,11) = 11.94, *p* < 0.01, η^2^_G_ = 0.42) and Tone (*F*(1,11) = 13.95, *p* < 0.01, η^2^_G_ = 0.30) and a significant interaction between them (*F*(1,11) = 5.72, *p* = 0.04, η^2^_G_ = 0.15). Bonferroni post-hoc test indicated that animals infused with saline showed increased freezing time during the post-tone period (*p* < 0.01). In contrast, animals infused with muscimol presented similar levels of freezing time before and after tone presentation (*p* = 1.00). Besides, the post-tone freezing time of the muscimol group was significantly lower than the saline group (*p* < 0.01). There was no difference between groups during the pre-tone period (first two minutes of the test, *p* = 0.98).

## 5. DISCUSSION

To the best of our knowledge, this is the first study investigating the specific roles of the aIC and the pIC in the consolidation of both CFC and TFC tasks. Our main finding is that post-training inactivation of the pIC, but not the aIC, selectively impaired conditioned freezing response to tone, suggesting that functional inactivation of the pIC during the consolidation time-window is sufficient to severely impair cued but not background contextual fear memories. These results are consistent with the growing evidence that the pIC plays an important role in emotional processing (Gogolla, 2017; Méndez-Ruette et al., 2019), and point to an important role of this insular area not only in integrating exteroceptive and interoceptive signals (Gehrlach et al., 2019; Livneh et al., 2020), but also in consolidating the association between neutral cues and an aversive exteroceptive stimulus. In addition, our results highlight the different functional profiles of aIC and pIC in fear memory consolidation.

Our findings corroborate those of Brunzell & Kim (2001), who showed a selective impairment of TFC after post-training electrolytic lesions of the pIC (Brunzell & Kim, 2001), however, a major limitation of their study was the probable damage of fibers of passage coursing through the lesioned area (Lavond et al., 1993). Our results, hence, support that the TFC impairment was not due to such damage or other nonspecific effects of such lesions. However, Corodimas & LeDoux (1995) reported that post-training lesions of an area (−1.80 to −3.80 mm from bregma) - partially corresponding to the here considered pIC - abolished the expression of freezing response to both tone and contextual conditioned stimuli (Corodimas & LeDoux, 1995). The CFC impairment in their study may be explained either by (a) potential damage of fibers of passage, as commented above; (b) the larger posterior extension of the lesion, encompassing the perirhinal cortex that is known to affect the conditioning freezing response to the context (Kent & Brown, 2012), and (c) the permanent character of the electrolytical lesions that may interfere with retrieval processes or freezing expression. Considering the evidence that post-training protein synthesis inhibition of the pIC immediately after, but not 6 hours following training, also impaired TFC (Casanova et al., 2016), our results clarify that the pIC is selectively recruited for the consolidation of cue-fear memories.

This selective role in memory consolidation of TFC is also supported by anatomical studies showing that the pIC is a candidate for the integration of viscerosensory inputs (Allen, 2020; Livneh et al., 2020) as well as for the convergence of both conditioned stimulus (CS, i.e., tone) and unconditioned Stimulus (US, i.e., footshock), resulting from multisensory afferences of thalamic nuclei, as well as from the lateral and basolateral nuclei of the amygdala (Gehrlach et al., 2020; McDonald & Jackson, 1987; Shi & Cassell, 1998a; Shi & Davis, 1999). Although there seems to be a consensus in the literature that the IC is not necessary for the acquisition of fear conditioning - since manipulations before training have shown no effect (Brunzell & Kim, 2001; Gehrlach et al., 2019; Lanuza et al., 2004) - it is possible that the pIC is preferably recruited to consolidate and maintain the association between a discrete stimulus (i.e., tone) and the US, which would explain the selective effect observed with post-training inactivation of the pIC in our study. On the other hand, our results also suggest that the formation or maintenance of a more complex association between context and footshock does not depend on either aIC or pIC.

The complete absence of effect with the post-training inactivation of the aIC is in contrast with previous studies showing that pre-training lesions or post-training neurotransmission inhibition of the aIC attenuated the conditioned freezing response to the training context (Alves et al., 2013; Morgan & LeDoux, 1995). It is of note, however, that the manipulated region in these studies comprised the more rostral area of the IC, which was not included in the present study. However, pre-training lesions of an area that corresponds to the aIC in our study are known to affect memory performance in tasks that require encoding of spatial or contextual components, such as the spatial version of the Morris water maze (Nerad et al., 1996) and the inhibitory avoidance task (Bermudez-Rattoni & McGaugh, 1991; but see Dunn & Everitt, 1988).

The lack of effect in CFC with post-training inactivation of the aIC in the present study may suggest that this IC subregion is only involved when the aversive stimulus is intense. Here we used a mild training protocol (single tone-footshock pairing) to avoid fear generalization during the TFC test, whereas Alves and colleagues (2013) employed a strong training session with 6 footshocks (1.5mA/3s), hence indicating that the aIC may be necessary for memory consolidation only after stronger CFC trainings (Alves et al., 2013). Accordingly, a growing body of evidence suggests that strong aversive events but not mild ones recruit the activation of the Salience Network, which comprises the aIC (Schwabe, 2017). In addition, the activation of glucocorticoid receptors in this region has been reported to enhance memory consolidation of both contextual and footshock components of the inhibitory avoidance task (Fornari, Wichmann, Atucha, et al., 2012). Thus, even though the aIC is not strictly necessary for CFC and TFC consolidation, the activation of this anterior IC subregion, either by stress hormones or higher training intensities, may modulate the strength of fear memories.

In conclusion, our study extends previous evidence about the role of IC in the emotional memory neural circuitry, showing functional differences along the rostral-caudal axis. While the aIC is not involved in memory consolidation for both CFC and TFC tasks, the pIC plays a selective role in the consolidation of cued-fear responses. Therefore, our results strengthen the importance to analyze the functional dissociation of the IC subregions in conditions related to fear memory such as post-traumatic stress disorder (Harricharan et al., 2020; Nicholson et al., 2016).

## ACKNOWLEDGEMENTS

This work was supported by *Universidade Federal do ABC* (UFABC), *Universidade Federal de São Paulo* (UNIFESP) and *Coordenação de Aperfeiçoamento de Pessoal de Nível Superior* (CAPES).

## Data

The data used for all analyzes in this manuscript can be found at https://data.mendeley.com/datasets/838krppypd/1

## Author contributions

JP: Investigation, Formal analyses, Writing - Original Draft, Project administration; AB: Investigation, Writing - Review & Editing the manuscript; MdSC: Formal analyses, Writing – Original Draft; TLF: Writing - Review & Editing the manuscript; MGMO: Resources, Writing - Review & Editing the manuscript; RVF: Conceptualization, Supervision, Writing - Review & Editing the manuscript. All authors approved the final version of the manuscript.

